# difFUBAR: Scalable Bayesian comparison of adaptive evolution

**DOI:** 10.1101/2025.05.19.654647

**Authors:** Hassan Sadiq, Patrick Truong, Maximilian Danielsson, Venkatesh Kumar, Hedwig Nora Nordlinder, Darren Patrick Martin, Ben Murrell

## Abstract

While many phylogenetic methods exist to characterize evolutionary pressure at individual codon sites, relatively few allow direct comparison between different *a priori* selected sets of branches. Such comparisons may be useful for pinpointing precisely the codon sites that are under differing selective pressures due to differing environmental contexts, or differing genomic contexts via epistatic interactions. Indeed, this was only recently addressed by an approach, developed in the frequentist framework, that proposes a site-wise likelihood ratio hypothesis test.

Previously, we have demonstrated that approximate grid-based Bayesian approaches to characterizing site-wise variation in selection parameters can outperform individual site-wise likelihood ratio tests. Such grid-based approaches can exhibit poor computational scaling when the number of site-wise parameters expands, but here we show that this is still tractable up to four parameters, and that a simple subtree-likelihood caching strategy can provide efficiency improvements in some cases.

We propose difFUBAR, which allows the demarcation of two branch sets of interest and, optionally, a background set, and estimates joint site-specific posterior distributions over *α, ω*_1_, *ω*_2_, and *ω*_*BG*_ using a Gibbs sampler. Evidence for hypotheses of interest can then be quantified directly from the posterior distribution, and we standardly report *P* (*ω*_1_ *> ω*_2_), *P* (*ω*_2_ *> ω*_1_), *P* (*ω*_1_ *>* 1), and *P* (*ω*_2_ *>* 1).

We characterize the computational and statistical performance of this approach on previous simulations, comparing it to the site-wise likelihood ratio test approach, where it shows moderate statistical benefits, and substantial computational gains, typically being more than two orders of magnitude faster on the same datasets. We also demonstrate that it can scale to datasets of over ten thousand taxa, on a laptop in under ten minutes. difFUBAR is implemented in `MolecularEvolution.jl` - a Julia framework for phylogenetic model development - and can be run locally, or online via a Colab notebook.

## Introduction

The structure of the genetic code, with synonymous and non-synonymous substitutions, leaves a footprint of adaptive evolution upon protein coding genes. Beginning with GY94^1^ and MG94^2^, various “codon model” approaches have been developed ^3^ ^4^ ^5^, which parameterize the substitution process over evolutionary trees in terms of an underlying nucleotide model, but with differential acceleration of non-synonymous and synonymous substitutions.

The primary aim of codon-substitution models is to detect departures from neutrality and to quantify the magnitude and nature of selection. Classic applications span diverse questions: likelihood-based models mapped clusters of positively selected residues in the HIV-1 *env* gene, illuminating immune-escape pathways ^6^; branch-site tests uncovered recurrent selection on primate TRIM5α, pinpointing the capsid-binding patch that governs species-specific retroviral restriction ^7^; and genome-wide codon analyses revealed adaptive change in oxidative-phosphorylation genes that met the extraordinary metabolic demands of powered flight in bats ^8^.

It is now well-established that both the underlying neutral substitution rate, and the strength of selection, can vary across sites ^9 10 11^. Approaches to model this site-wise variation can be broadly grouped into two categories: “random effects” models ^12 13 14^, that assume that the parameters of the codon model are drawn from a common distribution (which itself can be learnable), and perform inference using Bayes, or “empirical Bayes” methods ^15^, and “fixed effects” models ^16 17^, that assume distinct codon parameters for each site, and then perform site-wise likelihood hypothesis testing to answer statistical questions about the strength of selection at each site. Random effects models allow pooling of information across sites, which has statistical advantages, but original random effects models were computationally expensive, and could not scale to large datasets.

For codon models, there is a lack of closed-form tractability over the distribution of selection parameters, so random effects models rely on discretization schemes to approximate the distribution of selection parameters, which are represented by points on a parameter grid, and associated probability weights. Originally, the *α, β* values for these grid positions were themselves optimizable parameters, which meant that the total likelihood calculation scaled linearly with the number of grid points. This resulted in models with extremely coarse grid approximation schemes. In our previous work ^18^, we showed that, by fixing the grid values in advance and only learning the weights, we could re-cycle the likelihood calculations and provide a much finer-grained grid approximation (which ameliorates the fact that they are now fixed *a priori*) while simultaneously reducing the computational cost. This approach was implemented in FUBAR, which has subsequently become a popular tool for characterizing site-wise selection pressure.

In a different direction, Kosakovsky Pond et al. ^19^ developed Contrast-FEL, a site-wise likelihood ratio test to compare the strength of selection between two *a priori* defined sets of branches in a phylogeny. This approach allows a different kind of question to be asked of the sequence data, formulating the test not as a comparison of *ω* against the neutral expectation, but of *ω* between two (or more) sets of branches.

This can be applied to many evolutionary scenarios. For example, when a virus that typically circulates in bats infects and transmits in humans, a phylogeny constructed from sequences sampled from both species will have some branches representing within-bat viral evolution and some representing within-human viral evolution. A method that compares *ω* between these two sets of branches would identify codon sites that have been evolving under different selective constraints in humans vs in bats. This could, depending on the nature of the differences, provide clues into how the virus is adapting to its new host. Contrast-FEL, and its subsequent use, have clarified the value of such comparisons.

In this manuscript we propose difFUBAR, which aims to extend the grid-based Bayesian approach from FUBAR ^18^ to allow comparison of selection pressure between sets of branches, addressing the same kind of comparison as Contrast-FEL. difFUBAR specifically allows the comparison of selection pressure between two sets of branches, with optionally a background set of branches that do not participate in the comparison. Grid-based schemes can scale poorly with the number of parameters, but here we show that three and four parameters are well tolerated, and we also develop a subtree-likelihood caching strategy that can, for some trees, improve the efficiency of grid-based methods in this context.

difFUBAR is implemented in Julia ^20^, a language aimed at scientific computing, using the `MolecularEvolution.jl` phylogenetic modeling framework. We characterize the statistical performance of difFUBAR using previous simulations ^19^, comparing it to the Contrast-FEL site-wise likelihood ratio test approach, and demonstrate modest statistical benefits and dramatic improvements in efficiency and scalability.

## Methods

Like FUBAR, difFUBAR first computes a grid of conditional likelihood values for each site, where each grid point corresponds to a particular combination of parameter values. For difFUBAR, each branch uses an F3×4 codon model, parameterized with synonymous rate *α* and non-synonymous/synonymous rate ratio *ω* = *β/α*. Each branch can belong to up to three *a priori* designated branch classes: 1, 2, or *BG* (for “background”), which differ only by their value *ω*, with *ω*_1_, *ω*_2_ and *ω*_*BG*_, respectively (fig. 1A). Thus, the synonymous rate *α* varies from site to site, but is constant across all branches, and *ω* varies between the three branch classes.

**Fig. 1.**
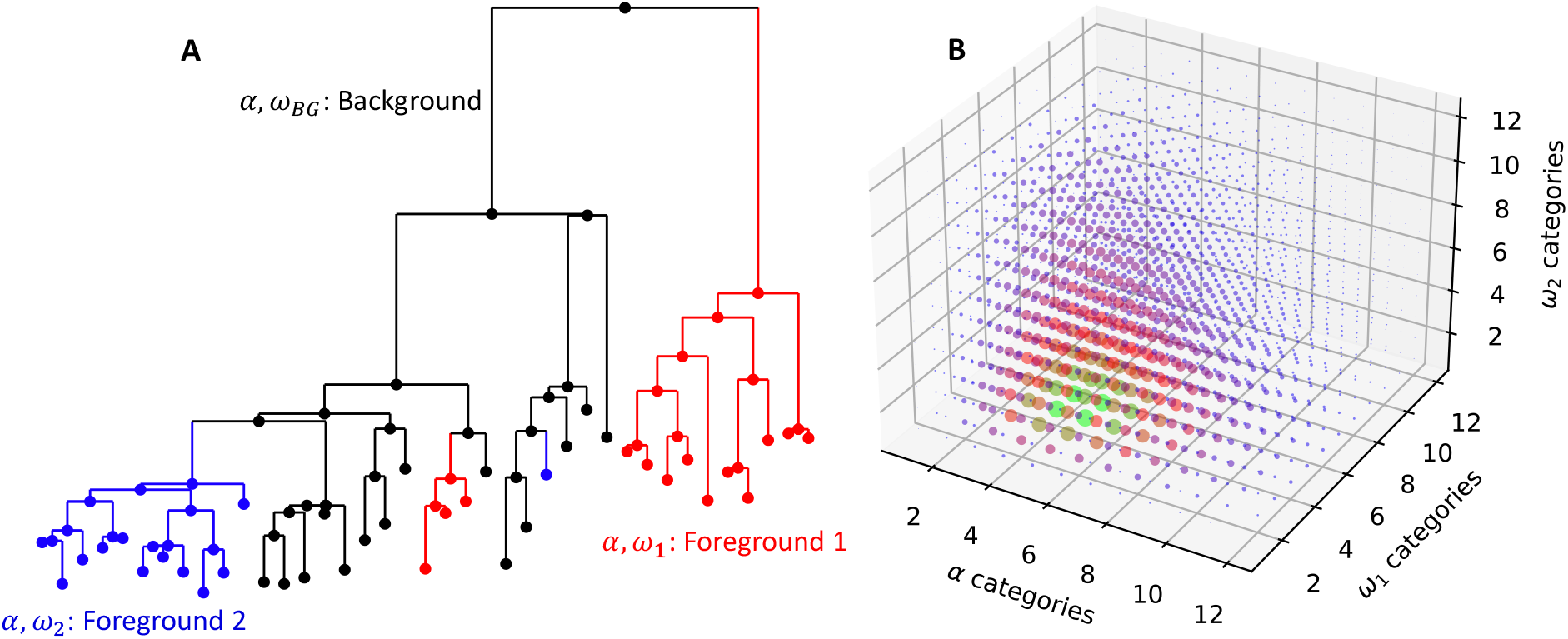
difFUBAR. difFUBAR allows the comparison, for individual codon sites, of selective pressure on different sets of branches on a tree. Panel **A** shows an example tree with two clades of interest: Foreground 1 and Foreground 2, and a set of Background branches that will not be involved in the comparison. Each branch set differs by selection parameter *ω*, which is the non-synonymous to synonymous substitution rate ratio. Thus each site has parameters *ω*_1_, *ω*_2_, and *ω*_*BG*_, as well as a synonymous rate parameter, *α* that is shared between all clades. We consider a discrete grid of parameter values, and any site can adopt any combination on the grid. Panel **B** shows an idealized example parameter grid (without *ω*_*BG*_ for ease of visualization), where each combination of parameters (ie. each point on the grid) is imbued with a weight, which is the probability that a site uses that combination of parameters. Evolutionary hypotheses (eg. *ω*_1_ *> ω*_2_) correspond to sets of points on the grid, and difFUBAR uses MCMC to marginalize over the alignment-wide category weights, computing site-specific posteriors for a pre-defined set of hypotheses. By default, difFUBAR reports posterior probabilities for four hypotheses – (a.) *ω*_1_ *> ω*_2_, (b.) *ω*_2_ *> ω*_1_, (c.) *ω*_1_ *>* 1 and (d.) *ω*_2_ *>* 1, examining positive selection on each branch-set of interest, as well as explicitly comparing the two sets of branches against each other.

For a site *i*, and a combination of our four codon model parameters, we can compute the probability of the site given the parameters (and implicitly conditioned on the phylogeny, and nucleotide model rates):

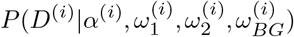

where *D*^(*i*)^ is the sequence data for site *i*. Our goal is then to infer, for all sites *i*,

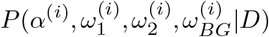

where *D* refers to the data at all sites.

To model evolution along a branch, *b*, of the phylogenetic tree, difFUBAR uses a continuous-time Markov process ^2^. The instantaneous rate matrix *Q* = *{q*_*ij*_*}* is defined as a 61 *×* 61 codon matrix (excluding the stop-codons) describing the rate of substitution of codon *i* with codon *j*:

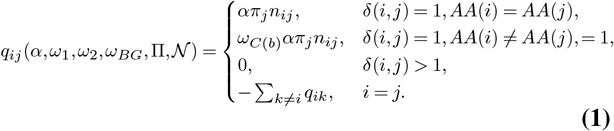

*δ*(*i, j*) denotes the number of nucleotide differences between codon *i* and *j, AA*(*i*) is the amino acid encoded by *i, C*(*b*) is the branch class which *b* belongs to, *n*_*ij*_ (comprising *N*) are the nucleotide mutational biases, which are modelled using a reversible GTR model ^21^ and *π*_*j*_ (comprising Π) are the equilibrium frequencies of the codons.

We estimate a global F3×4 codon model from the data, where *ω*_1_ = *ω*_2_ = *ω*_*BG*_ = *ω* = *β/α*, in two steps. First, we estimate *N* and branch lengths under a nucleotide model, and use the F3×4-estimator for Π. These parameters are then fixed to their estimates when we find the maximum likelihood estimates of *α* and *β*. We let 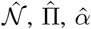 denote the estimates for *𝒩*, Π, *α* respectively. Then, branch lengths are scaled by a factor 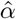 so that the maximum likelihood estimate of *α* is 1, which is the point we’ve chosen to center our grid approximation around. 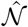 and 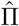are used throughout downstream.

Following FUBAR ^18^ we assume that the parameter values lie on fixed grid points (fig. 1B). By default, difFUBAR allows 12 grid values for *α, ω*_1_, and *ω*_2_, and 7 grid values for *ω*_*BG*_. Thus, when a tree has two groups of branches that are compared to each other, and a background set, there are 12, 096 *α, ω*_1_, *ω*_2_, and *ω*_*BG*_ parameter combinations (we call each combination a “category”, following the “Random Effects Likelihood” literature) whose likelihoods must be evaluated at each site. Without background branches, there are 1, 728 categories.

For each category, *C*, given a probability *θ*_*k*_ = *P* (*C*_*k*_), the marginal likelihood can be calculated as:

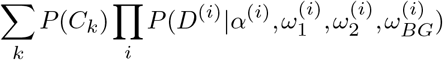

FUBAR used Metropolis-Hastings to sample from the posterior distribution of *θ* values, using a Dirichlet prior *P* (*θ*). Models with this structure have been well studied in the “Latent Dirichlet Allocation” ^22^ literature, and here we retain this model structure but replace FUBAR’s original Metropolis-Hastings approach with a much more efficient Gibbs sampler ^23^.

The Gibbs sampler iteratively samples the latent category allocations *z*_*i*_ for each site *i* and the category probabilities *θ* from their full conditional distributions. Using a symmetric Dirichlet prior with concentration parameter *γ* (default: 0.1), where *θ ∼* Dirichlet(*γ*):

1. For each site *i*, sample the category allocation *z*_*i*_ from:

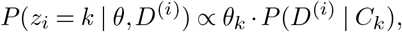

where *P* (*D*^(*i*)^ | *C*_*k*_) is the precomputed likelihood of site *i* under category *k*.
2. Sample the category probabilities *θ* from the Dirichlet posterior:

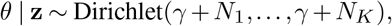

where 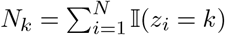 counts sites allocated to category *k*, and 𝕀 (*·*) is the indicator function.

After discarding burn-in iterations (default: 1/5 of the samples), the sampler produces draws from the joint posterior *P* (*θ*, **z** | *D*). Posterior probabilities for evolutionary hypotheses are approximated by the proportion of post-burn-in samples satisfying the hypothesis, which are accumulated during sampling.

Canonically we report posterior probabilities for 4 hypotheses of interest for all sites: *P* (*ω*_1_ *>ω*_2_), *P* (*ω*_2_ *>ω*_1_), *P* (*ω*_1_ *>* 1), *P* (*ω*_2_ *>* 1), but note that other measures of evidence (such as Bayes factors), or other hypotheses could also be reported with minimal modification.

In addition to posterior probabilities, we also offer visualization of the marginal distributions of sites identified under any hypothesis. These provide a visual explanation for cases where, for example, a site might have strong evidence for *ω*_1_ *>* 1, no evidence for *ω*_2_ *>* 1, but also no evidence for *ω*_1_ *≠ ω*_2_ (eg in fig. 3).

### Subtree-likelihood caching

For each combination of parameter values, difFUBAR would, naively, need a likelihood calculation for the entire tree, via Felsenstein’s algorithm ^24^. But given values of the four codon parameters, an intermediate step in computing the likelihood of *D*^(*i*)^ is to compute, for every node *s*, the joint probability of the unobserved codon state 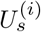 at that node and the observed data 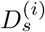 beneath it:

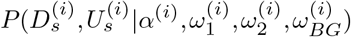

When a subtree rooted at node *s* is *pure*, consisting of descendant nodes that only belong to a single *ω*-group, *ω*_*k*_, the parameter values for the other *ω*-groups cannot affect it, and the above reduces to:

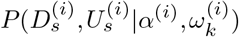

As illustrated in fig. 2, difFUBAR exploits this by finding *maximal pure* subtrees (a pure subtree where the parent of the root of the subtree is the root of a subtree that is not pure), and precomputing 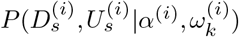 for these. These quantities are cached at the subtree root, and the sub-tree root’s descendants are pruned away, eliminating recalculation for all combinations of the *ω*-rates that do not affect the subtree probabilities. For trees with high *purity*, defined as the proportion of nodes that belong to a pure subtree (excluding the roots of maximal pure subtrees) to the total nodes in the tree, this can dramatically reduce the computational complexity of computing the grid of conditional likelihoods of all parameter combinations.

**Fig. 2.**
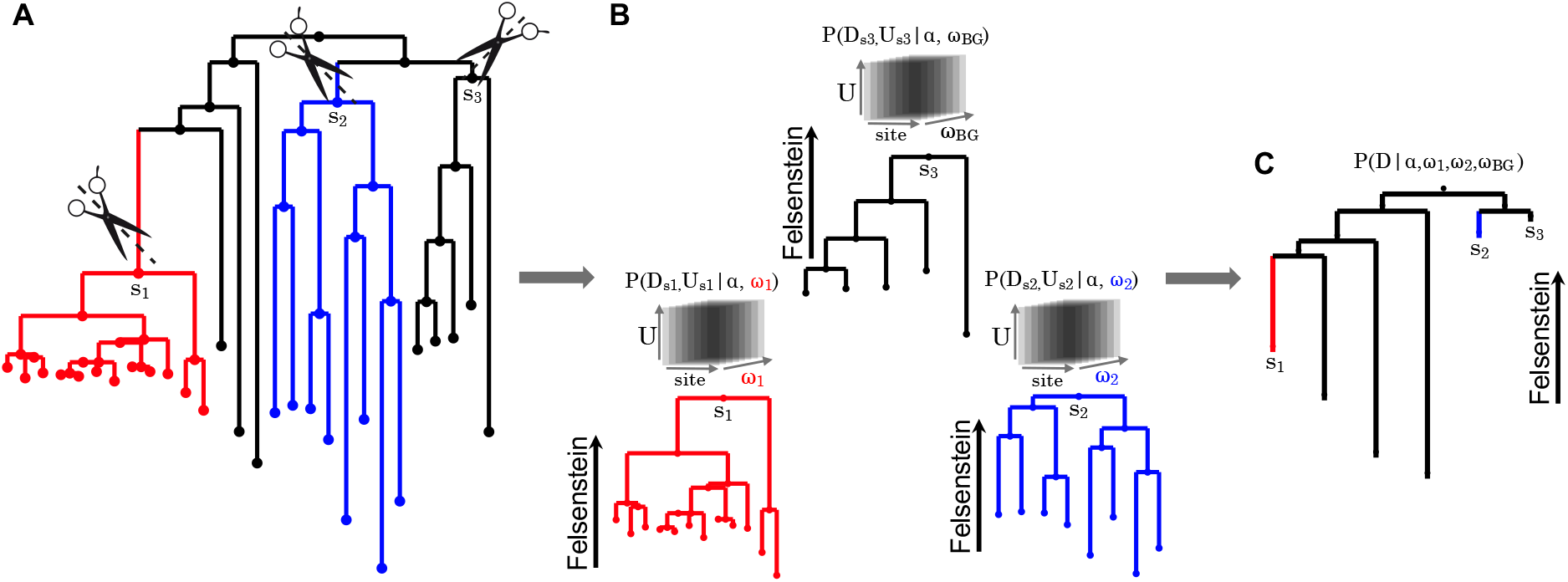
Subtree-likelihood caching. Where possible, we exploit the structure of the branch tags to avoid re-computing sub-tree likelihoods. Panel **A** shows an example tree where we have identified three pure subtrees, where all descendant nodes belong to the same *ω*-group. We disconnect the root nodes of these subtrees from the main tree, and (as shown in panel **B**) for each site we compute, for each pure subtree *s*, the joint likelihood of observing the sequences on the leaves of the subtree (*D*_*s*_) and the unobserved codon state (*U*_*s*_) at the subtree root: *P* (*D*_*s*_, *U*_*s*_|*α, ω*_*s*_), for all values of *ω*_*s*_, the only *ω* value used on all nodes beneath the subtree root. These codon state-by-site-by-*ω* likelihoods are cached and, as shown in Panel **C**, the subtree roots can be treated as leaves for the full Felsenstein likelihood calculation for all parameter combinations, retrieving the cached subtree likelihoods instead of recomputing them.

**Fig. 3.**
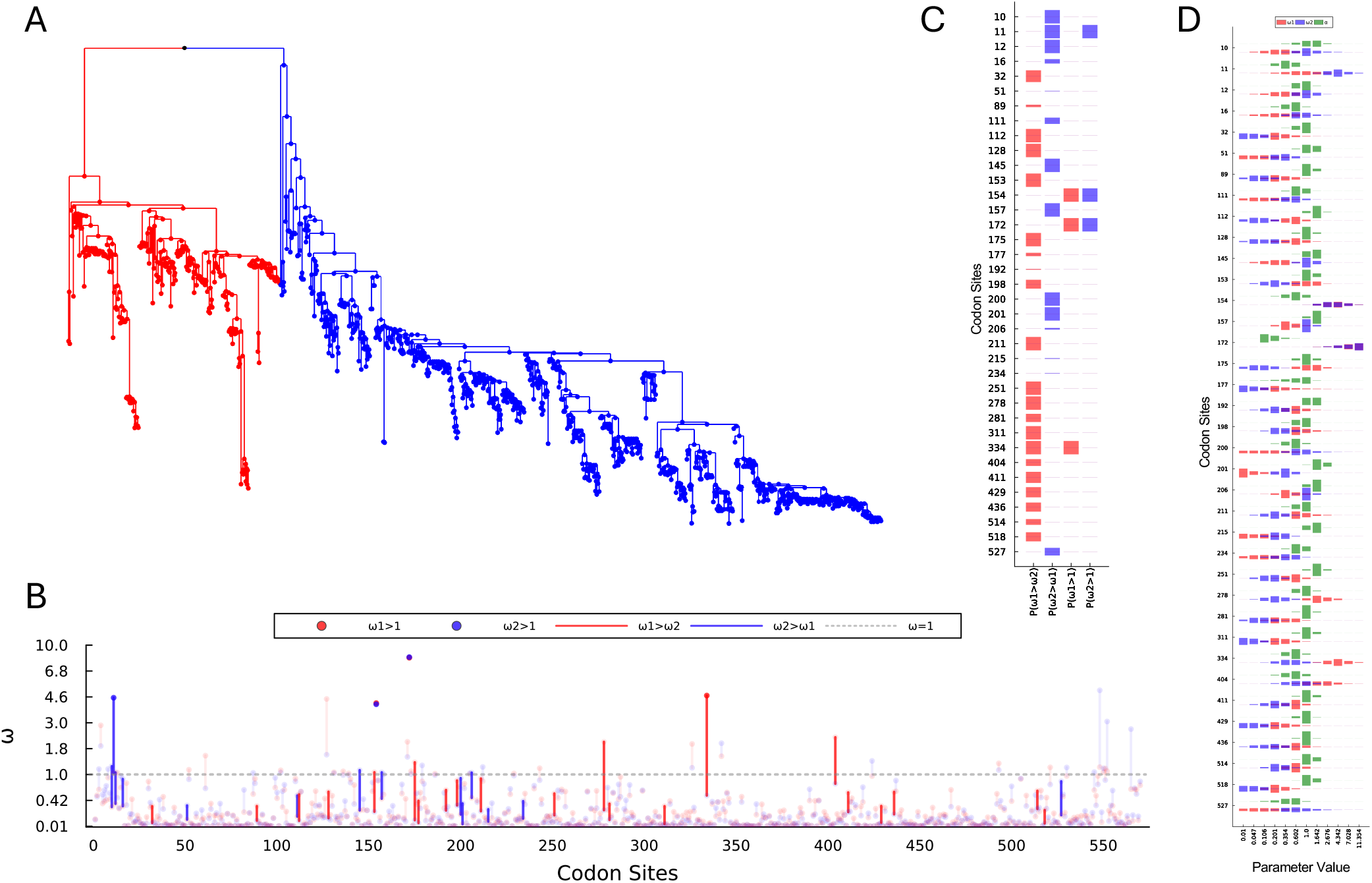
difFUBAR hypothesis visualization. difFUBAR outputs a standard set of visualizations, summarizing the complex space of hypotheses. As an example, Panel **A** shows a phylogeny of Influenza A H5 sequences, partitioned into two clades (red and blue). Panel **B** aims to provide an alignment-wide overview, while still depicting which hypotheses were above the chosen significance level (see legend). Panel **C** shows site-specific posterior probabilities for all four hypotheses, scaling bars between 0 (when the posterior was less than or equal to the threshold) to full size (as the posterior approaches 1). Panel **D** shows the marginal posterior distributions, for each site, over *α* and both *ω* parameters. This can be used to understand the signal driving significant differences (eg. large differences in parameters vs small differences but with high precision estimates).

### Method Availability

difFUBAR is implemented in `CodonMolecularEvolution.jl`, which is a codon model companion package to the `MolecularEvolution.jl` Julia framework. This framework provides, via Julia’s multiple dispatch, generic methods for phylogenetic likelihood calculations, branch length optimization, and tree topology search, as well as a host of convenience utilities.

Inspired both by the dramatic success of ColabFold ^25^ for making AlphaFold2 available to the wider biological community, as well as the recent availability of Julia kernels in Google Colaboratory, we developed a difFUBAR Colab note-book that allows one to upload one’s data and run difFUBAR in the cloud. This removes the need for local installation, and allows more customization than a typical webserver, since the analysis parameters, posterior threshold, etc can all be adjusted, and the plots customized.

difFUBAR takes standard alignment (.FASTA) and phylogeny (Newick) formats, but requires branches to be labeled. This can be done programmatically when the tagging strategy is such that the node labels can be inferred from the leaf names, or interactively via `phylotagger`, a web tool that allows branches to be selected and labeled, exporting the labeled tree in the correct format.

Resources can be found at:

- `CodonMolecularEvolution.jl`: https://github.com/MurrellGroup/CodonMolecularEvolution.jl
- `difFUBAR Colab notebook`: https://colab.research.google.com/github/MurrellGroup/ColabMolecularEvolution/blob/main/notebooks/difFUBAR.ipynb
- `Phylotagger`: https://murrellgroup.github.io/WebWidgets/phylotagger.html

For this manuscript, the analysis of simulation results was performed in Julia, and visualized using `Plots.jl` ^26^.

## Results

### Statistical Performance

To benchmark the statistical and computational performance of difFUBAR, we relied heavily on simulations established for Constrast-FEL in Kosakovsky Pond et al. ^19^, available at https://data.hyphy.org/web/contrast-FEL/simulations/. Constrast-FEL is parameterized with a synonymous rate *α*, and non-synonymous *β* parameters for each group of branches, and we consider the case with two sets, such that we have *β*_1_ and *β*_2_. The *α, β* (Constrast-FEL) vs *α, ω* (difFUBAR) parameterization choice can affect inference, but is not a meaningful difference for simulations.

We use the “omnibus” simulation set, which provides i) phylogenies ii) two labeled branch sets for each phylogeny, iii) simulation parameter settings, specifying the *α, β*_1_ and *β*_2_ values for each codon site, iv) fasta formatted alignments of sequences simulated over the phylogenies with these parameters, and v) Contrast-FEL results. This comprises 530 phylogenies and parameters, with five replicates for each.

For tests that compare *β*_1_ to *β*_2_, which include Contrast-FEL’s likelihood ratio test, and two of difFUBAR’s reported posterior probabilities, sites simulated under the null hypothesis of *β*_1_ = *β*_2_ provide a means to benchmark the false positive rates of these methods. Further, the simulations encompass a wide range of values, and it is expected that the power to detect differences in selection will increase, all else being equal, for more divergent *β*_1_ and *β*_2_ values. We thus stratify our analyses into ranges of |*β*_1_ *− β*_2_|, which is used as a measure of the effect size.

By design, difFUBAR’s tests that compare *ω* parameters on different branch sets are directional (ie. either *ω*_1_ *> ω*_2_ or *ω*_2_ *> ω*_1_). For the purposes of this benchmarking study, since the Constrast-FEL analyses provide a single p-value, we consider difFUBAR to have detected a difference between the two clades if the posterior for either directional hypothesis was above the threshold (equivalently, we use the maximum of the two directional posterior probabilities).

For the results depicted in Fig. 4, we pool all sites from all “omnibus” simulations, and group them into effect size ranges. For each effect size, Receiver Operator Characteristic (ROC) curves are computed using the sites within that effect size window, combined with all the null sites (which are required to quantify the false positive rate). Fig. 4 A shows that difFUBAR and Contrast-FEL have similar power for very low effect sizes (where both methods are performing just above random), which is not unexpected for data generated by a stochastic substitution process. However, at larger effect sizes difFUBAR appears to have a notable advantage, with moderately superior power for the same false positive rates.

**Fig. 4.**
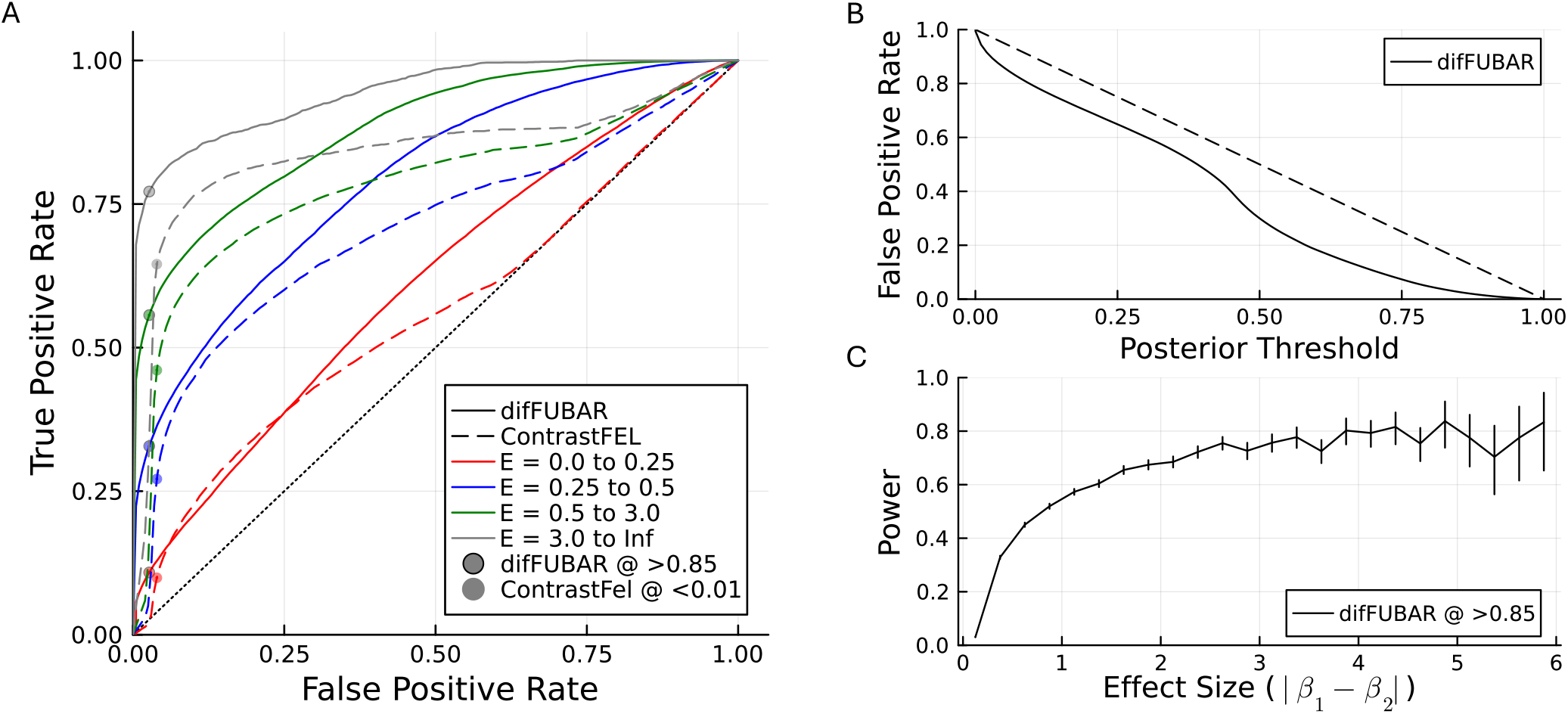
Statistical performance for tests comparing. *β*_1_ **to** *β*_2_ was assessed using simulations from Kosakovsky Pond et al. ^19^ (the “omnibus” set), with data from 530 alignments of varying numbers of sequence and codon sites, with *α* varying over sites, and *β* varying over sites and, sometimes, between two sets of *a priori* identified branches. For this comparison, the null hypothesis is true where *β*_1_ = *β*_2_, and we use |*β*_1_ *− β*_2_ | as a measure of the effect size. difFUBAR provides directional posteriors (ie. either *ω*_1_ *> ω*_2_ or *ω*_2_ *> ω*_1_), but we wish to compare to to ContrastFEL’s single p-value, so we take the maximum of the two directional posteriors for this purpose. Panel **A** shows Receiver Operator Characteristic (ROC) curves, examining the false positive and true positive rates for all posterior (or p-value) thresholds, split into effect size ranges. We annotate thresholds of *p* = 0.01 for Constrast-FEL, and posterior *>* 0.85 for difFUBAR. Panel **B** shows how difFUBAR’s false positive rate varies with the posterior threshold, for this set of simulated data. Panel **C**, with the posterior threshold fixed at 0.85, shows how power increases with effect size (with binomial Clopper-Pearson 95% confidence intervals).

**Fig. 5.**
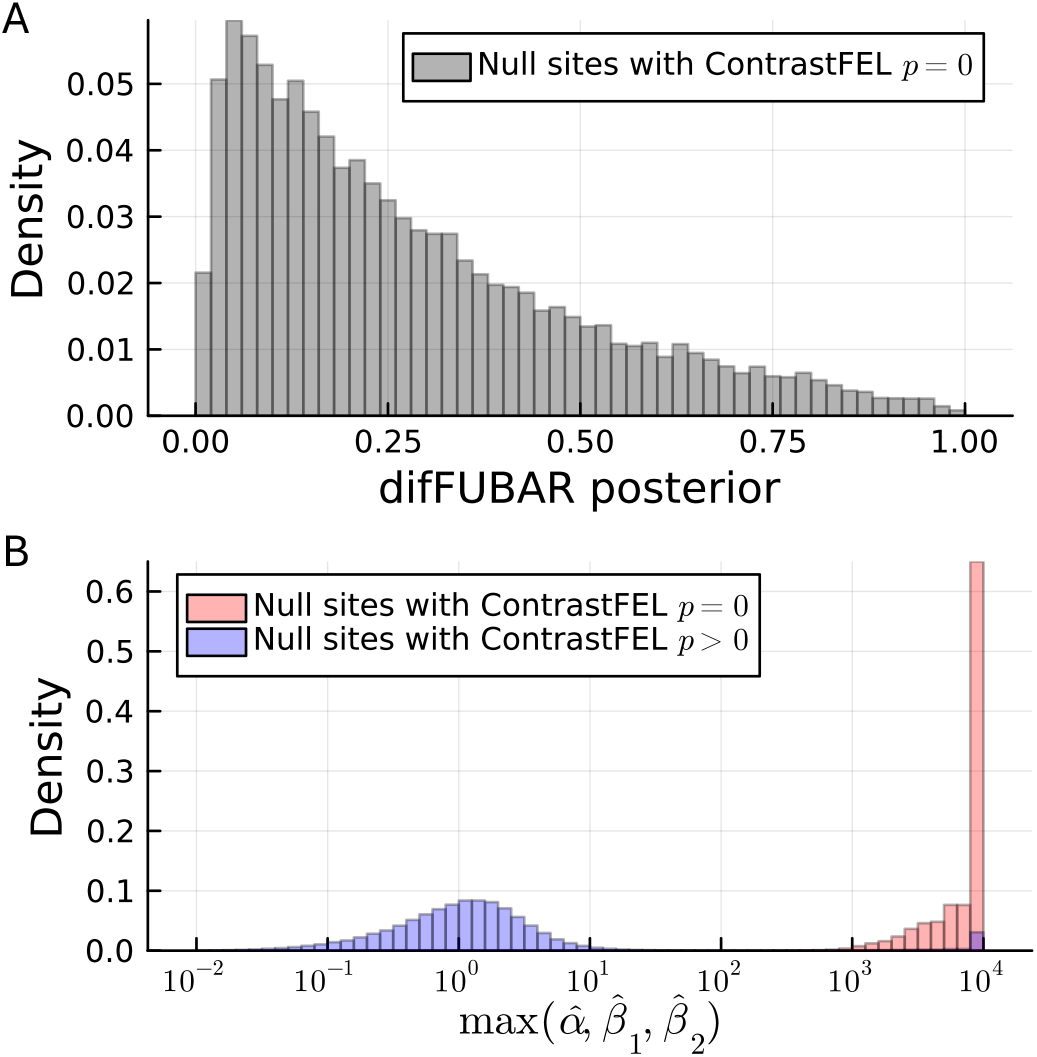
Frequentist false positives. From the “Omnibus” simulation analysis outputs provided with the Contrast-FEL publication, a subset of sites simulated under the null hypothesis had extremely low Contrast-FEL p-values (*≈* 0). Panel *A* shows that, per difFUBAR’s posterior (the max of *P* (*ω*_1_ *> ω*_2_) and *P* (*ω*_2_ *> ω*_1_)), these did not generally show strong evidence of differences between the clades. Panel *B* shows that, for these sites, Contrast-FEL’s point estimates of either *α, β*_1_ or *β*_2_ were numerically extreme, which was not typically the case for null sites with non-zero p-values. This supports the hypothesis that these extreme p-values may be driven by difficulties with the site-wise likelihood maximization required for Contrast-FEL’s p-value calculations. Note that a preliminary investigation suggests that this issue may have been remedied in more recent versions of Contrast-FEL.

As Bayesian posterior probabilities do not necessarily have frequentist coverage properties, simulations can also be useful for investigating the behavior of the detection thresholds. Fig. 4 A shows that a posterior threshold of 0.85 appears to control the false positive rate relatively well, while also being well powered to detect sites that actually differ. Fig. 4 B shows that, as the posterior threshold approaches 1, the false positive rate drops dramatically, with extremely low false positive rates for thresholds of 0.9 and above in these simulations. Further, Fig. 4 C shows that, with this threshold, the power climbs steadily with the |*β*_1_ *− β*_2_| effect size, approaching 80% power for the larger range of effect sizes explored in these simulations.

### Extreme p-values

We noticed that a proportion of sites simulated with *β*_1_ = *β*_2_ (ie. null sites) had Contrast-FEL p-values of *≈* 0, sufficiently common to create a visible effect in the bottom left of the Contrast-FEL ROC curves (4 A). We used difFUBAR, as an independent method, to evaluate whether these null sites were somehow exhibiting anomalous evidence of between-branch selection pressure differences by chance. The maximum of difFUBAR’s two directional *ω*_1_ *> ω*_2_ or *ω*_2_ *> ω*_1_ posteriors skews low, however, showing no indication that these sites exhibit any evidence of selection differences (5 A). Further investigation identified that, at almost every single one of these sites, the Contrast-FEL maximum-likelihood parameter estimates for either *α, β*_1_, or *β*_2_ are numerically extreme, as shown in 5 B where the max of these three values is in the 1000+ range, whereas for null sites with *p >* 0 the corresponding distribution was mostly in a reasonable range. This suggests the possibility that these extreme p-values, inferred from null sites, are driven by optimization difficulties during site-wise likelihood maximization.

### Efficiency and Scalability

As with the original FUBAR, compute efficiency, and the corresponding data size scalability that this purchases, was a key motivating factor in the development of difFUBAR. Using a relatively standard workstation configuration, with an Intel(R) Core(TM) i9-10980XE CPU @ 3.00GHz 64-bit architecture with 36 CPUs, we benchmarked the runtime of difFUBAR on each of the datasets in the “omnibus” simulation. difFUBAR was launched using `julia -threads 20`, using just over half of the workstation compute.

By default, difFUBAR uses a heuristic that depends in part on the *ω*-group purity, to choose between pure subtree likelihood caching, and more aggressive parallelization, which can take advantage of more CPUs without subtree caching. The branch labeling exhibited in the “omnibus” simulations tends toward lower degrees of *ω*-group purity, with an average of only 1/4 of the nodes falling within a pure clade, which reduces the benefit of difFUBAR’s subtree likelihood caching. As shown in Fig 6 A, difFUBAR is extremely fast, never taking longer than 30 seconds on any of the “omnibus” simulations. Further, difFUBAR’s runtime is reliably linear in both the number of codon sites, and the number of sequences, in line with it’s theoretical complexity. Some degree of efficiency gain is exhibited with higher tree purity, but this is very minimal, likely due to the more moderate tree purities, and the availability of 20 CPUs which affords reasonable efficiency for non-subtree-caching parallelization strategies. On the same hardware, we also ran Contrast-FEL (HYPHY v2.5.72) on a random subset (N=58) of these simulations, using MPI for parallelization, and using all 36 available CPUs. For the datasets tested, difFUBAR was typically around two orders of magnitude faster than Contrast-FEL, and this difference seemingly increased for larger datasets, sometimes approaching three orders of magnitude faster.

**Fig. 6.**
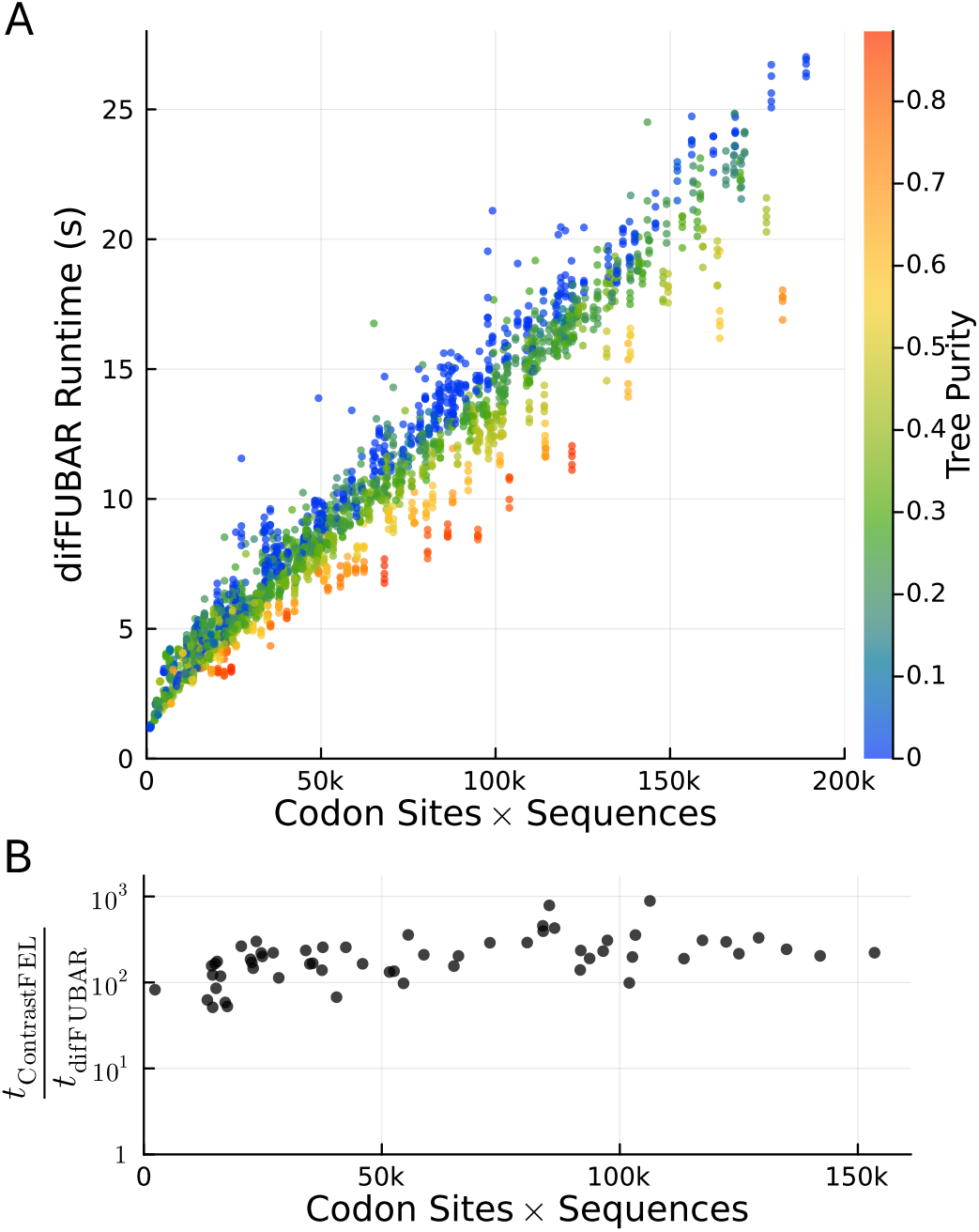
Execution time. Using simulated data from Kosakovsky Pond et al. ^19^, Panel **A** shows that difFUBAR is, above some minimum, predictably linear in the number of codon sites and the number of sequences being analyzed. Further, it is faster for higher tree purity values (as expected from the subtree likelihood caching technique). For a randomly sampled subset of datasets, run on the same hardware, Panel **B** shows that difFUBAR is typically two (and sometimes nearly three) orders of magnitude faster than Contrast-FEL, where the relative speedup appears to increase with size of the input data.

More targeted investigation showed that subtree likelihood caching can make a large difference. On an Apple Silicon M3 Max laptop, the analysis in Fig 3, which has 679 sequences and 570 alignment sites, and which has a branch labeling that allows that maximum exploitation of subtree likelihood caching, takes 27 seconds. If one instead labels leaf nodes chaotically, removing the subtree caching benefit, it takes 161 seconds - around six-fold slower.

To investigate whether these efficiency gains would permit the analysis of substantially larger datasets, we constructed a version of the H5 dataset in 3, but using NCBI Virus (https://www.ncbi.nlm.nih.gov/labs/virus/) to obtain all H5 sequences. After filtering, and de-duplication, there were 11,547 sequences, and the resulting codon alignment had 588 codon sites. Using a tagging strategy as in 3, tagging the same two major clades, the entire analysis took just under 10 minutes on an M3 Max laptop.

## Discussion

difFUBAR extends the grid-based conditional likelihood pre-computation strategy, initially developed for FUBAR, to the problem of comparing selective pressure between sets of branches on a phylogenetic tree. Grid-based approaches scale exponentially poorly in the number of parameters that must be captured by the grid. While FUBAR sought to model just two parameters varying over sites (*α* and *β*), difFUBAR requires three (*α, ω*_1_, *ω*_2_) in the ‘no-background’ case, and a fourth (*ω*_*BG*_) if there are background branches. As a result, instead of FUBAR’s 400 grid points, with difFUBAR’s default discretization it has 1,728 or 12,096 parameter combinations that must be evaluated for each site.

Despite this, our results show that difFUBAR’s advantages over Contrast-FEL are very much in line with FUBAR’s over FEL. Like FUBAR, it has a modest improvement in power for a given false positive rate, likely driven by the implicit regularization of Bayesian inference, and its ability to weakly share distributional signals across sites. Also like FUBAR, which exhibited a large runtime advantage over FEL’s site-wise likelihood ratio test, difFUBAR shows a dramatic efficiency advantage over Contrast-FEL’s sitewise testing approach. This might seem unexpected given the exponential increase in grid points as the number of parameters increases, but that neglects that the site-wise likelihood ratio testing approaches have to solve higher dimensional maximization problems for each site as the grid size increases. Empirically, for this many parameters, grid approaches are still performant.

Our exploration of the extreme Contrast-FEL p-values in the context of difFUBAR’s posterior evidence suggests that this statistical issue was driven by optimization difficulties, which is always a concern when a method must solve multiple maximization problems for each site. If optimization under the null or alternative hypothesis gets stuck in a local optimum, or if there are any other optimization difficulties, then the p-values could be extremely wrong in either direction. Rather than being an idiosyncrasy of this particular comparison, however, this is an inherent difference in robustness between grid-based Bayesian methods and site-wise frequentist methods. The Gibbs sampler used to infer the alignment-wide grid weight parameters mixes extremely quickly, and at the level of individual sites, all grid points are explicitly summed over. This makes it impossible to fail to explore a region of the parameter space, so long as it is encompassed by the grid. We would like to note, however, that a preliminary investigation of the small Omnibus subset used here for timing comparisons suggests that this issue may have been improved in more recent versions of Contrast-FEL.

One common statistical error is to test a hypothesis about two quantities, and over-interpret a difference in the outcome of the two tests. For example, one might test for positive selection at a site on two different clades, and find that *ω*_1_ is significantly greater than 1, but *ω*_2_ is not significantly greater than 1, and mistakenly conclude that there is statistical support for a difference between *ω*_1_ and *ω*_2_. This is nicely encapsulated in the title of the classic Gelman and Stern ^27^ ‘The Difference Between “Significant” and “Not Significant” is not Itself Statistically Significant’. Part of the motivation behind difFUBAR itself, but especially behind some of the visualizations (eg. in 3), is to make it entirely clear when differences in *ω* are significant, rather than differences in significance. One limitation of this approach is that, unlike Contrast-FEL, difFUBAR cannot easily be extended to test for any differences between more than two clades. Statistical power can be gained by conducting a single hypothesis test using multiple groups at once, pooling evidence from all groups without diluting it via multiple pairwise comparisons. Further work will be required to explore the statistical properties of using difFUBAR in a pairwise fashion in such scenarios.

One area of possible development is the Dirichlet prior over the grid weights. While inferred grid weights (eg. posterior means) for these methods are typically smooth, this smoothness is due to an effect of averaging, and is not encoded in the prior, which treats each grid point independently. Are there statistical advantages to be unearthed by encoding, in the prior, the expectation that neighbouring grid points should have more similar weights? And does this issue become more important for difFUBAR’s extremely large grids compared to FUBAR’s relatively small ones? Approaching this will likely require developing efficient sampling strategies for priors that lack any form of tractable conjugacy.

The demonstration that grid-based approaches comfortably expand to three and four parameters raises the prospect that they might be useful for a wider family of methods in this space. Approaches that allow variation over time, either encoded in the CTMC (eg. Covarion codon models ^28^) or via branch-wise random effects (eg. BS-REL ^29^ and MEME ^30^) often have to handle site-level variation as well, and may benefit from FUBAR-like grid-based approaches.

## ACKNOWLEDGEMENTS

This project has received funding from the Swedish Research Council (2022-05034) to B.M.

## AUTHOR CONTRIBUTIONS

Conceptualization, H.S., V.K., B.M.; Formal Analysis, H.S., P.T., M.D., V.K., H.N.N., B.M.; Investigation, H.S., P.T., M.D., H.N.N., B.M.; Resources, M.D., V.K., B.M.; Visualization, H.S., P.T., M.D., H.N.N., B.M.; Writing – Original Draft, H.S., P.T., M.D., B.M.; Writing – Review & Editing, All authors; Funding Acquisition, B.M.

